# Discovery of allele-specific protein-RNA interactions in human transcriptomes

**DOI:** 10.1101/389205

**Authors:** Emad Bahrami-Samani, Yi Xing

**Author notes:** Corresponding author: Yi Xing, Department of Microbiology, Immunology, and Molecular Genetics, University of California, Los Angeles, Los Angeles, CA 90095-7278.

## Abstract

Gene expression is tightly regulated at the post-transcriptional level through splicing, transport, translation, and decay. RNA-binding proteins (RBPs) play key roles in post-transcriptional gene regulation, and genetic variants that alter RBP-RNA interactions can affect gene products and functions. We developed a computational method ASPRIN (Allele-Specific Protein-RNA Interaction), that uses a joint analysis of CLIP-seq (cross-linking and immunoprecipitation followed by high-throughput sequencing) and RNA-seq data to identify genetic variants that alter RBP-RNA interactions by directly observing the allelic preference of RBP from CLIP-seq experiments as compared to RNA-seq. We used ASPRIN to systematically analyze CLIP-seq and RNA-seq data for 166 RBPs in two ENCODE (Encyclopedia of DNA Elements) cell lines. ASPRIN identified genetic variants that alter RBP-RNA interactions by modifying RBP binding motifs within RNA. Moreover, through an integrative ASPRIN analysis with population-scale RNA-seq data, we showed that ASPRIN can help reveal potential causal variants that affect alternative splicing via allele-specific protein-RNA interactions.

---

Natural genetic polymorphisms can diversify the transcriptome and proteome among individuals by altering the post-transcriptional processing and modification of RNA [1]. Such regulatory variation can cause disease, modify disease risk, or affect therapeutic response [2, 3]. Thus, the discovery of genetic variants that affect post-transcriptional RNA regulation may reveal causal mechanisms underlying phenotypic variability and disease pathogenesis in human populations [4, 5].

RNA-binding proteins (RBPs) are key regulators of post-transcriptional RNA processing and modification [6]. RBPs participate in various steps of RNA regulation, including splicing, transport, translation, and decay, thus determining the fate of RNAs after transcription [7]. RBPs bind to their RNA targets via defined sequence and/or structural motifs [8].

The predominant technology for transcriptome-wide mapping of RBP-RNA interactions is CLIP-seq [9–12]. Multiple variants of CLIP-seq (HITS-CLIP [9], PAR-CLIP [10], iCLIP [11], and eCLIP [12]) aimed at improving library efficiency and reducing artifacts, have been used to define the RBP-RNA binding landscape of hundreds of RBPs across different cell types and species. These variants of CLIP experiment are all fairly similar in essence, which is cross-linking RBP and its targets for a more stringent washing of unbound RNA followed by high-throughput sequencing, but due to their technical differences and biases, deliver slightly different datasets, as detailed in [13].

Previous studies have investigated the effects of genetic variants on post-transcriptional regulation, primarily using a sequence motif-based approach. Jian *et al*. [14] reviewed eight bioinformatics tools that predict splice-altering single nucleotide variants in the human genome. These methods use information about highly conserved splicing regulatory elements (5’ and 3’ splice sites and branch point signals) as well as auxiliary *cis*-acting elements recognized by trans-acting RBPs [14] to predict the effects of genetic variants on alternative splicing. Some other recent studies used defined binding motifs of RBPs to predict variants that alter RBP-RNA interactions [15, 16]. However, as RBP binding motifs are typically short (4-6 nucleotides) and degenerate, methods based on RBP motifs are expected to have a low accuracy and high noise [17].

We developed ASPRIN (Allele-Specific Protein-RNA Interaction), a computational method to identify genetic variants that alter RBP-RNA interactions via a joint analysis of CLIP-seq and RNA-seq data. The premise of ASPRIN is that the allelic ratio in CLIP-seq data compared to that in RNA-seq data of the same cell type can reflect the effects of genetic variants on RBP-RNA interactions. We performed a systematic ASPRIN analysis of CLIP-seq and RNA-seq data for 166 RBPs in two ENCODE cell lines. ASPRIN identified genetic variants that alter RBP-RNA interactions by modifying conserved RBP binding sites. Moreover, through an integrative ASPRIN analysis with population-scale RNA-seq data, we showed that ASPRIN can help reveal causal variants that affect alternative splicing via allele-specific protein-RNA interactions. The ASPRIN source code and user documentation are freely available for download at: https://github.com/Xinglab/ASPRIN.

## Results

### ASPRIN pipeline for detecting allele-specific protein-RNA interactions

The discovery of allele-specific protein-RNA interactions in ASPRIN is based on the rationale that if a particular single nucleotide polymorphism (SNP) creates or disrupts an RBP binding site, we would expect to observe a difference in the allelic ratio of the SNP in the CLIP-seq reads compared to the corresponding RNA-seq reads from the same cell type. A schematic diagram of ASPRIN is provided in Fig. 1a. Briefly, to call SNPs, RNA-seq reads were mapped to the human genome and transcriptome, and single nucleotide variants (SNVs) were called using the GATK pipeline [18] (see details in Methods). We then applied stringent filters to remove false positive SNPs contributed by potential sequencing errors, alignment artifacts, and RNA editing events. Specifically, heterozygous variants in RNA-seq data that matched known SNPs in the NCBI SNP Database (dbSNP) were kept [19], while potential RNA editing events were removed by intersection with the RADAR (rigorously annotated database of A-to-I RNA editing) RNA editing database [20]. After this set of high-confidence SNPs was generated, CLIP-seq reads were mapped and reads supporting the reference or alternative allele in the CLIP-seq data were counted. Additionally, because RNA-seq reads are typically longer than CLIP-seq reads, we split the 100 bp RNA-seq reads in the ENCODE data into two 50 bp segments and mapped them separately to count reference and alternative alleles in the RNA-seq data, to alleviate systematic mapping bias for the reference over the alternative alleles in CLIP-seq data compared to the RNA-seq data. This procedure largely removed the bias for reference over derived alleles in the ASPRIN results (Supplementary Fig. 1). Finally, we tested each SNP site with at least ten reads (sum of two alleles) in both the RNA-seq and CLIP-seq data for significant difference in allelic ratio via Fisher’s exact test of allelic read counts in RNA-seq versus CLIP-seq data. After correcting for multiple hypothesis testing, we reported SNPs with corrected *p*-values of less than 0.1 as ASPRIN SNPs (Fig. 1a). An example result for the HepG2 cell line is an A-to-G SNP (rs115776575) in the *PTPN4* gene that disrupts a highly conserved “A” nucleotide in the “TGCATG” consensus motif of RBFOX2. While the allelic ratio between “A” and “G” was 1:1 in the RNA-seq reads, the “G” allele represented only 10.5% of the CLIP-seq reads (Fig. 1b), consistent with RBFOX2 binding to the TGCATG motif, and that the A-to-G SNP at the fourth nucleotide position of the motif disrupts RBFOX2 binding.

**Figure 1:**
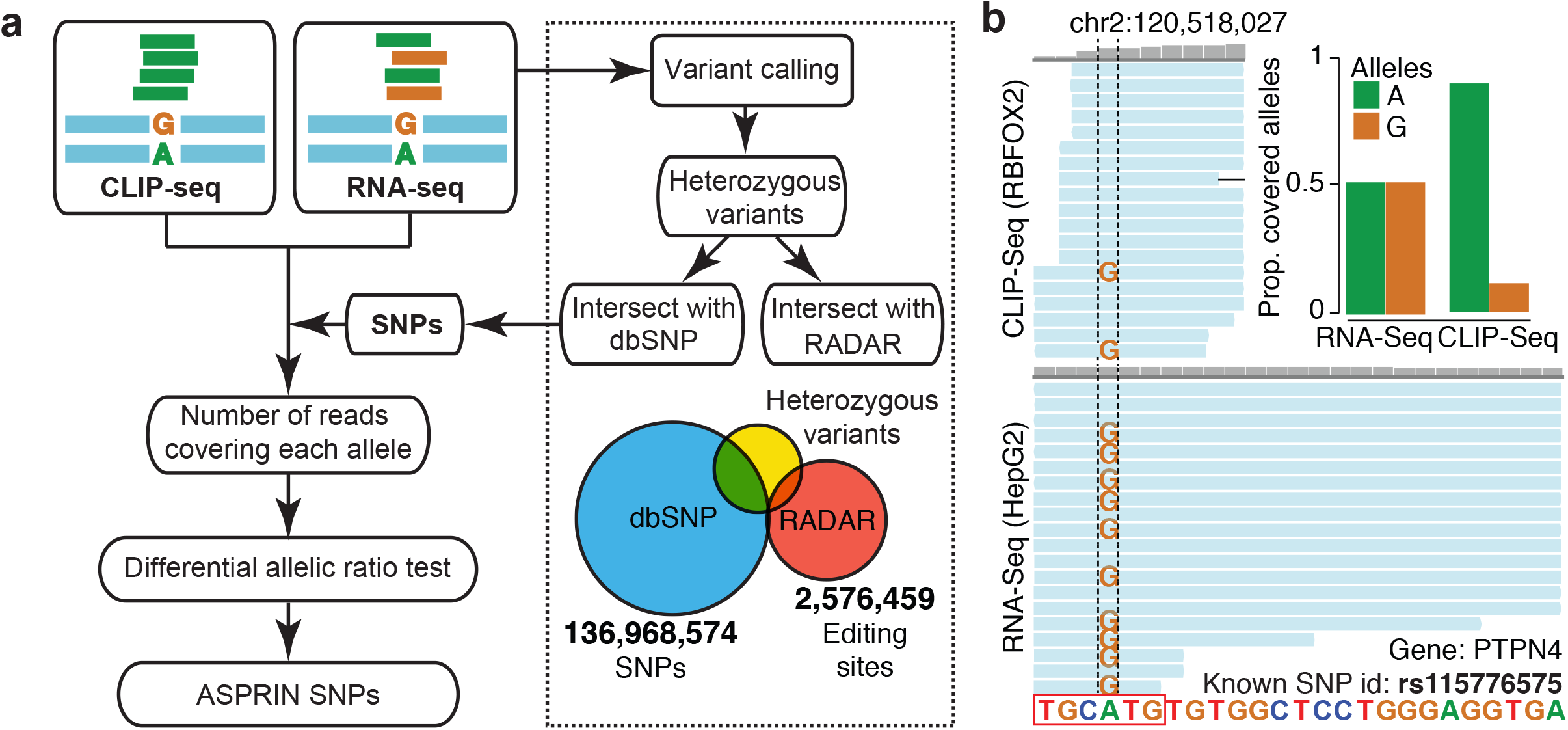
The ASPRIN pipeline for identifying allele-specific protein-RNA interactions from CLIP-seq and RNA-seq data. (**a**) Flowchart of the ASPRIN pipeline: variants are called from RNA-seq data, and heterozygous variants are intersected with dbSNP to obtain a list of high-confidence SNPs and intersected with RADAR to filter out potential A-to-I RNA editing events. For each SNP, ASPRIN counts the number of reads in the CLIP-seq and RNA-seq data that support each allele. An allelic ratio test then assesses whether one allele is significantly more preferred for RBP binding. (**b**) An A-to-G SNP (rs115776575) disrupts a consensus RBFOX2 binding site in the *PTPN4* gene. This disruption of binding is illustrated in the difference in the numbers of reads containing each allele in CLIP-seq reads, while equal numbers of reads contain each allele in the RNA-seq data.

### ASPRIN is robust in discovering SNPs involved in allele-specific protein-RNA interactions

We evaluated various issues that may affect the performance of ASPRIN, such as errors arising from calling variants from RNA-seq data, choice of RNA-seq protocols, and potential artifacts due to cross-linking step in CLIP-seq experiments. First, since whole-genome genotype data is not available for most of the cell types with CLIP-seq data, we assessed our SNP calling procedure using RNA-seq data alone. To obtain a ground truth for this assessment, we called SNVs using RNA-seq data for the GM12878 cell line (SRA accessions: SRR307897 and SRR307898), for which high-quality whole genome genotype data is available from the 1000 Genomes (1000G) Project [21]. After calling SNVs in GM12878 using our pipeline, we intersected the set of heterozygous variants with known SNPs in GM12878 from the 1000G project [21] and known A-to-I RNA editing sites in the RADAR database [20] to investigate the distribution of different variant types. As shown in Fig. 2a, 63.2% of the called SNVs were known SNPs, and 23.8% were known RNA editing events. The remaining 13.0% were unknown variants that did not match any 1000G SNPs or RADAR sites, and the distribution of all 12 possible single nucleotide changes suggested that these unknown variants represented a mixture of SNPs and RNA editing events (Fig. 2a). As shown in Fig. 2b, 89.6% of the called SNVs that were in the dbSNP were also present in the 1000G data for GM12878, suggesting an upper bound of 10.4% for the false discovery rate of our RNA-seq-based SNP calling procedure. Moreover, 3.6% of the called SNVs for GM12878 were present in the 1000G data but not in the dbSNP, suggesting that the use of the dbSNP had a minimal impact on the false negative rate of SNP identification. Collectively, our data suggest that, by using dbSNP and RADAR as filters, we can obtain a set of high-confidence SNPs from our RNA-seq variant calling in the absence of matching genotype data.

**Figure 2:**
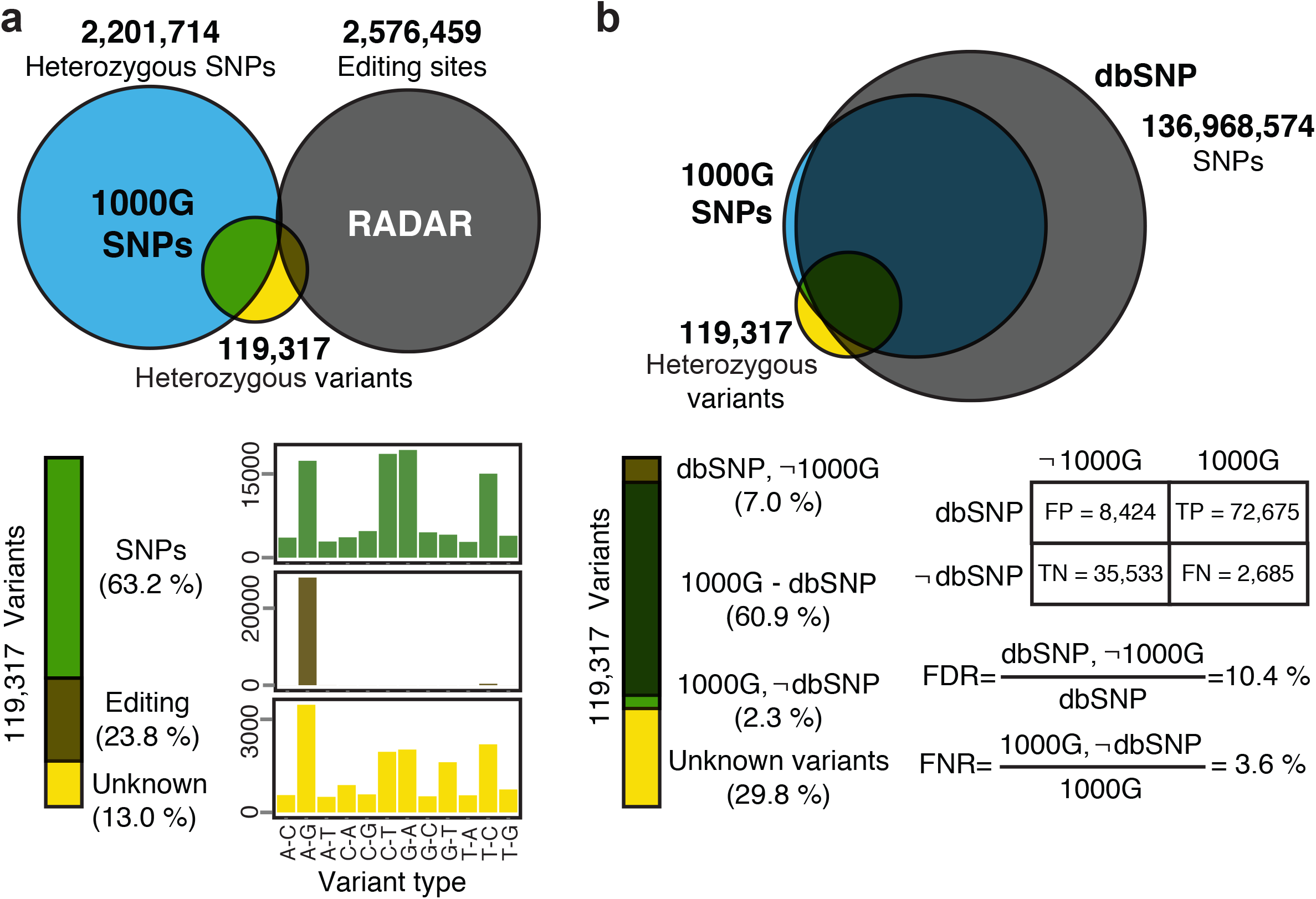
RNA-seq variants called in the GM12878 cell line. dbSNP and RADAR were used as external references to obtain a set of high-confidence SNPs from the RNA-seq variant calling in the absence of matching genotype data. (**a**) Intersection of variants with the 1000G SNPs and RADAR RNA editing events as well as the distributions of variant types over all 12 possible single nucleotide changes. (**b**) The variant filtering steps in the ASPRIN pipeline yields low false discovery and low false negative rates.

Next, we investigated issues that may affect the power of ASPRIN for calling SNPs and identifying allele-specific protein-RNA interactions. Specifically, the choice of RNA-seq protocol may affect the power of ASPRIN depending on the binding location of a given RBP within the RNA. For instance, a cytosolic polyA+ RNA-seq library would be appropriate for an RBP that predominantly binds to exons within mRNAs in the cytosol, but not for an RBP that predominantly binds to introns within precursor mRNAs in the nucleus. To investigate the most appropriate RNA-seq protocols and libraries, we randomly sampled equal numbers of reads from polyA+ and total RNA-seq libraries of distinct subcellular fractions (nucleus, cytosol, and whole-cell) from the HepG2 cell line and performed SNP calling and ASPRIN analysis on the sampled RNA-seq data. For both polyA+ and total RNA-seq libraries, we called the highest number of SNPs from the nuclear RNA-seq data and the lowest number of SNPs from the cytosolic RNA-seq data (Fig. 3a). The lowest number of SNPs was called from cytosolic polyA+ RNA-seq data (Fig. 3a); these SNPs were enriched for exonic regions within UTRs (Untranslated Regions) and CDS (Coding Segments) and depleted for intronic regions within pre-mRNAs (Fig. 3b). A similar trend was observed for the K562 leukemia cell line (Supplementary Fig. 2). On the other hand, as reads of cytosolic polyA+ RNA-seq libraries were concentrated within CDS and UTR regions, such data may have better power for detecting allele-specific protein-RNA interactions of RBPs that bind predominantly to exons. As expected, the nuclear fraction of the total RNA-seq library provided a much greater power for ASPRIN analysis of an RBP that binds predominantly to introns (HNRNPM), while ASPRIN analyses of an RBP that binds predominantly to exonic regions (YBX3) identified similar numbers of ASPRIN SNPs from the cytosolic polyA+ RNA-seq library and the nuclear total RNA-seq library (Fig. 3c). Furthermore, after calling peaks, we sorted all RBPs in both cell lines based on the ratio of exonic (CDS and UTR regions) to intronic peaks. The complete distributions of peaks in different regions for all RBPs are shown in Supplementary Fig. 3 and we excluded RBPs for which more than 50% of peaks fell in intergenic and noncoding regions. We observed a positive correlation (Pearson correlation coefficient = 0.34, *p*-value<0.0001) between binding of an RBP to exonic regions and the relative power of identifying significant ASPRIN SNPs using cytosolic polyA+ RNA-seq libraries, despite large variation among individual RBPs (Fig. 3d).

**Figure 3:**
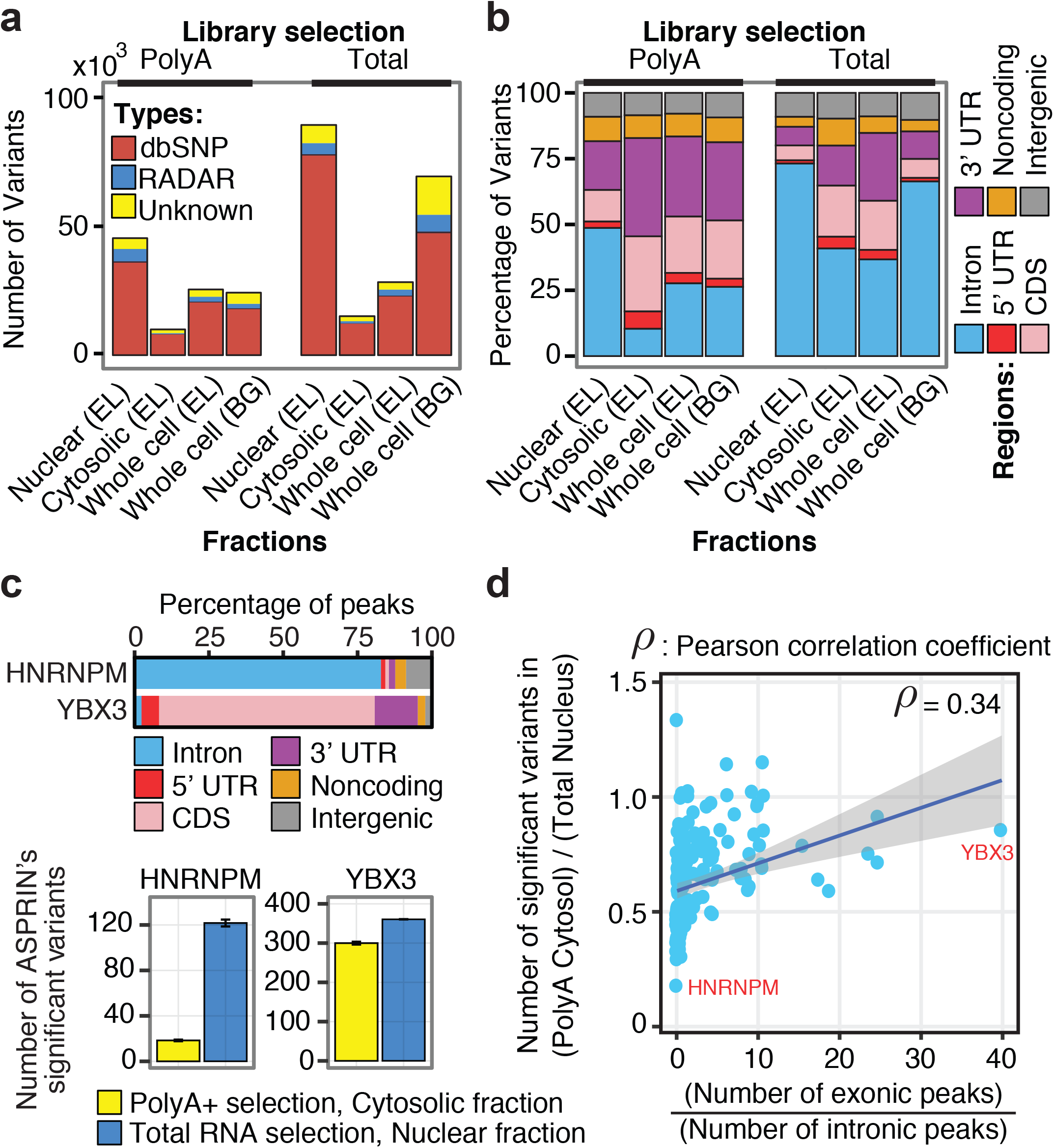
RNA-seq variants called from different RNA-seq libraries of the HepG2 cell line. Two methods of library selection (polyA+ and total RNA) in different subcellular fractions (nucleus, cytosol, and whole-cell fractions from two different labs: EL=Eric Lecuyer’s lab at Institut de Recherches Cliniques de Montréal, and BG=Brenton Graveley’s lab at University of Connecticut). (**a**) Numbers of variants called from different RNA-seq libraries and their intersections with dbSNP and RADAR. (**b**) Distribution of called variants in different genomic regions. (**c**) Numbers of significant ASPRIN variants from polyA+ cytosolic or total RNA nuclear RNA-seq libraries for an RBP that binds predominantly to intronic regions (HNRNPM) and an RBP that binds predominantly to exonic regions (YBX3). (**d**) The ratio of ASPRIN SNPs found using polyA+ cytosolic RNA-seq libraries to ASPRIN SNPs found using total RNA, nuclear RNA-seq libraries increases as the ratio of exonic to intronic peaks increases.

Finally, we evaluated potential false positives that may arise from the cross-linking step in CLIP-seq experiments. Specifically, the sequences in the CLIP-seq libraries may be altered by mutation or deletion at the cross-linking site [9–11]. We noted that in the eCLIP protocol used for generating the ENCODE CLIP-seq data, the majority of fragments were truncated at the cross-linking site rather than containing mutations or deletions [12]. Nonetheless, we investigated this issue further by calling SNVs from the ENCODE eCLIP data and comparing the distribution of variant types to that of the RNA-seq data and observed a similar distribution (Supplementary Fig. 4). Another possible source of artifacts is cross-linking bias that may shift the read count towards specific nucleotides in the CLIP-seq data. However, 70% of ASPRIN SNPs were called significant for only one RBP. Only 6% of ASPRIN SNPs were called significant for more than five RBPs. Among these SNPs, the same allele was preferred by all RBPs in 87% of the SNPs, whereas in the remaining 13% different alleles were preferred by different RBPs (Supplementary Fig. 5). Overall, these data suggest that the fraction of ASPRIN SNPs that may be attributable to CLIP-seq cross-linking bias is small.

### ASPRIN identifies functionally relevant SNPs for different classes of RBPs

To assess the potential functional relevance of the ASPRIN results, we investigated the positional distribution of ASPRIN SNPs for different classes of RBPs. To this end, we classified RBPs based on their known functions [22], and we defined genomic regions as follows: (1) 5’ UTRs; (2) upstream proximal intronic regions (500 nucleotides upstream of an internal exon); (3) coding regions; (4) downstream proximal intronic regions (500 nucleotides downstream of an internal exon); (5) 3’ UTRs; (6) distal intronic regions (more than 500 nucleotides away from exons on both sides); (7) noncoding regions; and (8) intergenic regions. Then, for each RBP, we calculated the enrichment of ASPRIN SNPs in different genomic regions (see details in Methods). As expected, ASPRIN SNPs were more enriched in regions to which RBPs bind to perform their known functions (Fig. 4). For instance, in the HepG2 cell line, we observed an enrichment (*p*-value < 0.001) of ASPRIN SNPs in the 5’ UTR for translation regulators such as DDX3X and NCBP2, with 27.1% and 16.0% of their ASPRIN SNPs found within the 5’ UTR respectively. Multiple classes of splicing factors showed distinct patterns of positional distributions for their ASPRIN SNPs. We observed an enrichment of ASPRIN SNPs in upstream proximal intronic regions for branch point recognition factors such as SF3B4 (30.5%), U2AF2 (12.2%), U2AF1 (8.8%), and SF3A3 (11.8%). Similarly, ASPRIN SNPs were enriched in the downstream proximal intronic regions for RBPs that are part of the 5’ splice site machinery such as PRPF8 (22.0%), EFTUD2 (14.8%), and RBM22 (11.7%). There was an enrichment of ASPRIN SNPs in coding regions for several splicing regulators that primarily bind to coding exons, such as SRSF1 (40.7%) and TRA2A (29.0%). For RBFOX2, we observed an enrichment of ASPRIN SNPs in both upstream and downstream proximal intronic regions (6.0% and 15.1%, respectively), as we expect RBFOX2 to bind to either region to promote exon skipping or inclusion respectively. The ASPRIN SNPs of HNRNP proteins were enriched in distal intronic regions and depleted in coding regions, which fits that these RBPs predominantly bind to distal intronic regions. Finally, RBPs that regulate mRNA stability, such as IGF2BP proteins and LIN28 showed an enrichment of ASPRIN SNPs in the 3’ UTR (Fig. 4a). We observed a similar pattern in the K562 cell line (Fig. 4b). The numbers of ASPRIN SNPs for each RBP in HepG2 and K562 are provided in Supplementary Fig. 6.

**Figure 4:**
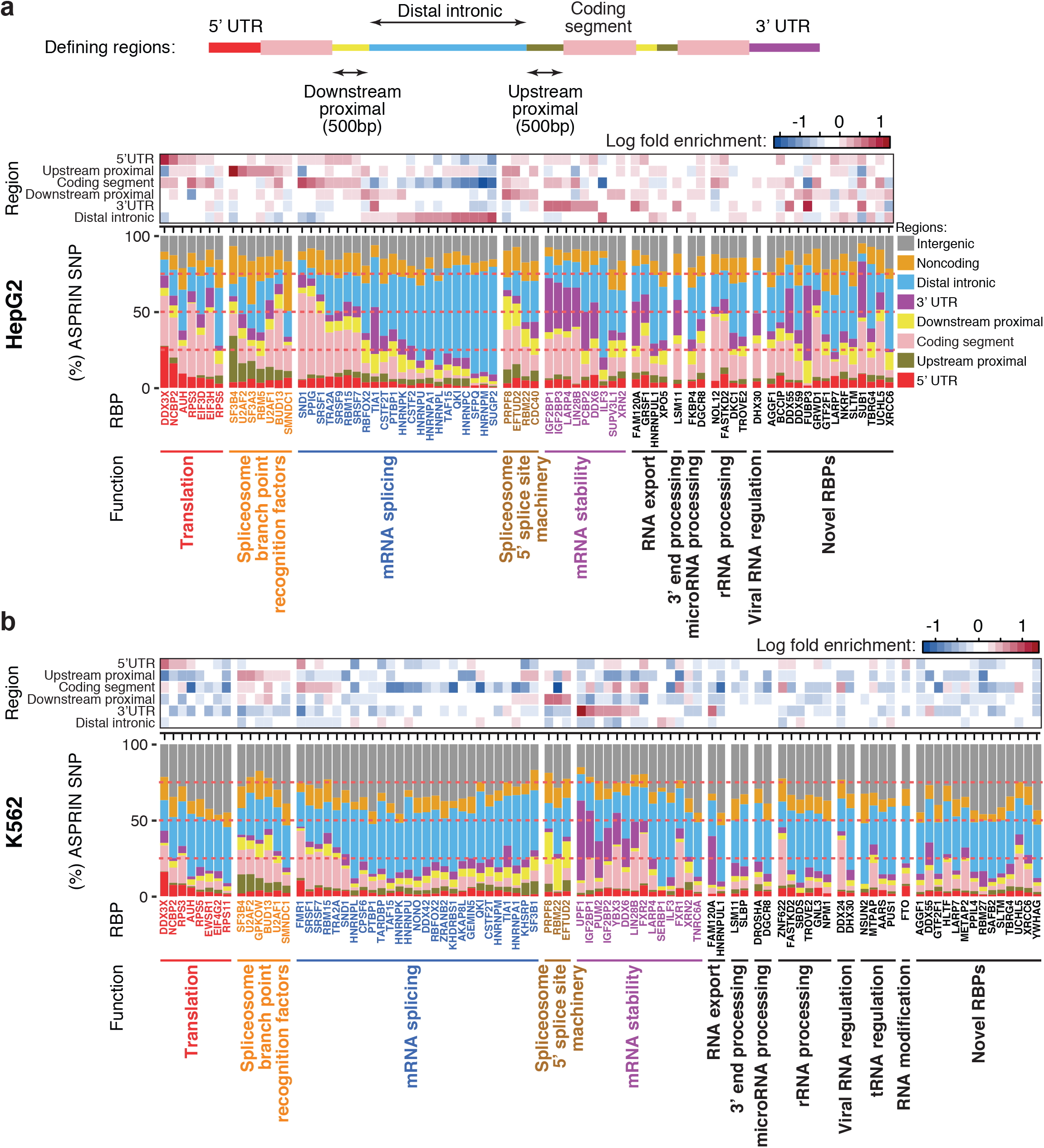
Enrichment of ASPRIN SNPs in different genomic regions. Positional distributions of ASPRIN SNPs for different classes of RBPs in HepG2 (**a**) and K562 (**b**) cell lines. The top diagram of the Figure depicts the different genomic regions used in the analysis. RBPs were classified based on their known functions [22]. In both panels the enrichment of ASPRIN SNPs for each RBP in different genomic regions is shown as heatmaps for color coded log fold enrichment (top) and barplots for % of total ASPRIN SNPs (bottom).

### ASPRIN SNPs affect RBP consensus motifs

To explore the potential molecular mechanisms by which ASPRIN SNPs affect protein-RNA interactions, we investigated the effects of ASPRIN SNPs on RBP consensus motifs. We predicted that if an RBP binds to RNAs in a highly sequence-specific manner, then variants within the conserved RBP consensus motif are likely to affect binding. First, we called peaks from ENCODE CLIP-seq data using Piranha [23] and performed *de novo* motif discovery on called peaks using Zagros [8] to obtain a 6-nucleotide consensus motif for each RBP. We then calculated the information content of the consensus motif, defined as the average information content of each position within the 6-nucleotide motif, as a measure of sequence specificity (see details in Methods). Fig. 5a shows the RBPs in HepG2, sorted by the sequence specificity of their consensus motifs. Among all RBPs, HNRNPA1 and RBFOX2 had the highest sequence specificity of their consensus motifs, and they are known to bind to highly conserved AGGGAG [24] and TGCATG [25] motifs, respectively. Next, for all ASPRIN SNPs of a given RBP, we obtained two sets of sequences that corresponded to the two alleles, i.e., one with high binding affinity and the other with low binding affinity. Finally, we used the position weight matrix that was obtained for all RBP consensus motifs by Zagros and calculated the motif scores for the two sets of sequences using STORM [26] (Supplementary Fig. 7 and Methods). Fig. 5b shows the motif scores of five RBPs with high (HNRNPA1, RBFOX2), median (DKC1), and low (NCBP2, XRN2) consensus motif sequence specificity. Variants in different positions within the consensus motif did not seem to affect binding equally. For example, for HNRNPA1, variants in position 5 of the motif had a more significant effect on binding than did variants in other positions. This result shows that not all positions in the consensus motif contribute equally to RBP-RNA interactions.

**Figure 5:**
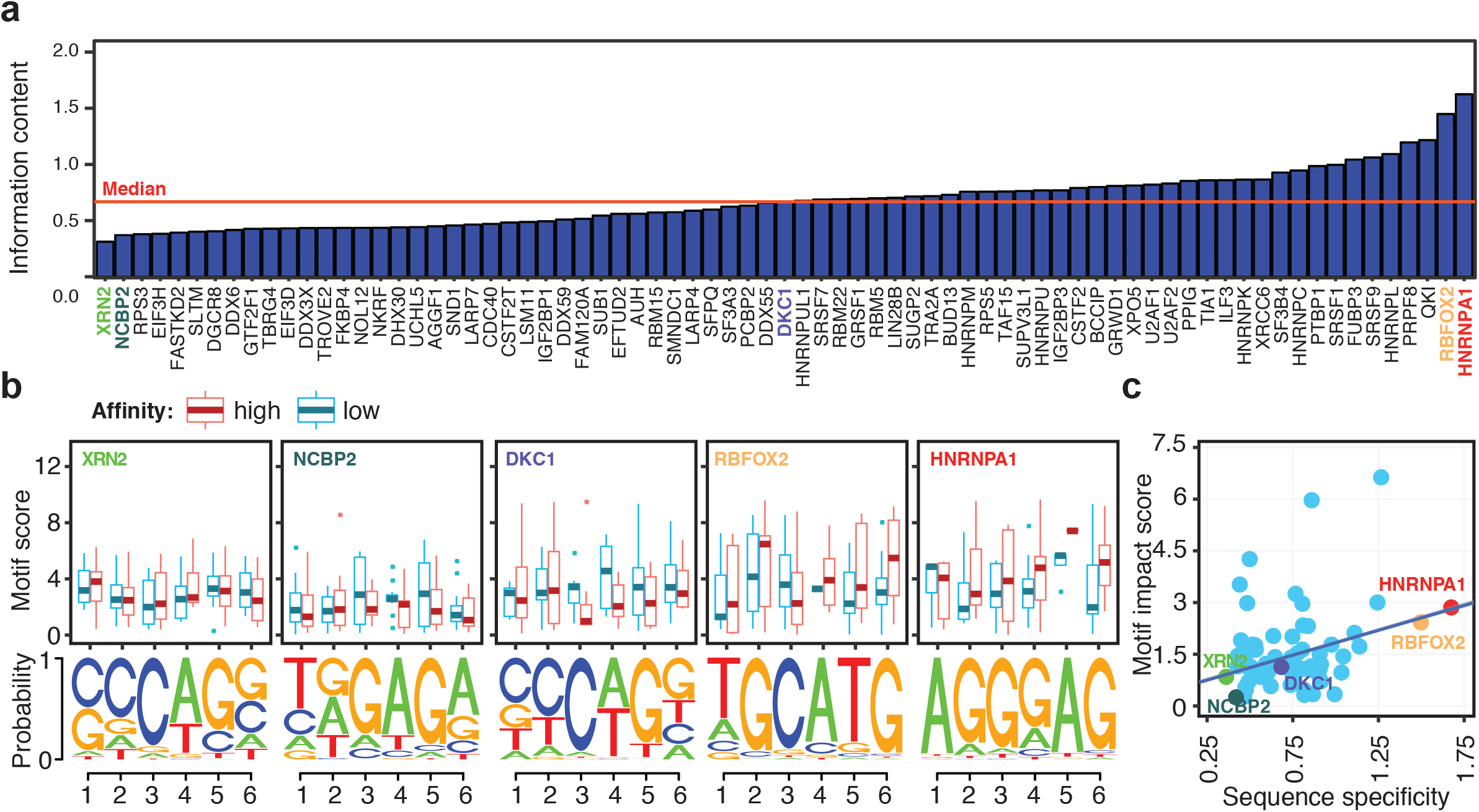
The effect of ASPRIN SNPs on RBP consensus motifs. (**a**) RBPs in the HepG2 cell line, sorted based on the sequence specificity (i.e. information content) of their consensus motif. For each RBP, the information content was calculated by taking the average of the information content for each position within the motif, calculated using Shannon’s entropy. (**b**) Boxplots comparing the consensus motif scores for alleles with higher and lower binding affinity. Two RBPs with the lowest sequence specificity (XRN2 and NCBP2), one RBP with the median sequence specificity (DKC1), and two RBPs with the highest sequence specificity (RBFOX2 and HNRNPA1) are shown. The consensus motif obtained from the top 1000 peaks for each RBP is represented at the bottom of each graph. (**c**) As sequence specificity increases, we observe a higher difference between the consensus motif scores of the high affinity versus low affinity ASPRIN alleles.

To further explore the relationship between the ASPRIN SNPs and RBP consensus motifs, we defined a motif impact score for each RBP and its associated ASPRIN SNP set as the maximum difference of average motif score between the two alleles with high versus low binding affinity in the window of 6 nucleotides overlapping the ASPRIN SNP (see details in Supplementary Fig. 7). We observed a positive correlation (Pearson correlation coefficient = 0.29, *p*-value<0.05) between the motif impact score and the sequence specificity of a given RBP’s consensus motif (Fig. 5c), suggesting that for highly sequence-specific RBPs, ASPRIN SNPs tend to affect binding by altering the consensus binding motifs within the RNA. For instance, in case of HNRNPA1 and RBFOX2, we observed a higher motif score for alleles with higher binding affinity, while for NCBP2 and XRN2, we did not observe noticeable differences in motif scores between the two alleles in any position of their consensus motif (Fig. 5c).

### ASPRIN can help reveal causal variants affecting alternative splicing

Finally, we investigated whether ASPRIN can help reveal causal genetic variants that affect post-transcriptional gene regulation. For this analysis, we focused on the genetic variation of alternative splicing. A series of population-scale transcriptome studies have revealed widespread alternative splicing variation among human individuals [4], but it remains challenging to pinpoint the causal genetic variants underlying this splicing variation. To match our ASPRIN analysis of the HepG2 liver cell line, we analyzed liver RNA-seq data along with matching genotype data of 71 individuals from the GTEx consortium (v6). We performed a transcriptome-wide scan of splicing quantitative trait loci (sQTLs) using GLiMMPS [27] and obtained ASPRIN SNPs correlated with GLiMMPS sQTLs (see details in Methods).

Our joint ASPRIN and GLiMMPS analyses revealed candidate causal SNPs that affected alternative splicing via allele-specific protein-RNA interactions. For example, GLiMMPS identified several SNPs that were significantly associated with an exon-skipping event in FAM114A1, one of which was an ASPRIN SNP (Fig. 6a). The genotype at the ASPRIN SNP was significantly associated with the level of exon inclusion, with the GG and AA genotypes showing the highest and lowest levels of exon inclusion, respectively (Fig. 6b). The ASPRIN analysis indicated that the G allele was associated with significantly greater binding by the splicing factor SRSF9 (Fig. 6b), while the A allele disrupted binding at this highly conserved “G” nucleotide at the fourth position of the SRSF9 consensus motif (Fig. 6b). Collectively, these data suggest that the G-to-A SNP disrupted the binding of the splicing activator SRSF9, leading to reduced inclusion of the FAM114A1 exon. Similarly, we identified an ASPRIN SNP for the splicing factor SF3B4, which was significantly associated with an alternative 3’ splice site event in ARL6IP4 (Fig. 6c, d). This C-to-T SNP was located 7 nucleotides upstream of the intron-exon boundary and disrupted a highly conserved ‘C’ nucleotide at the fourth position of the SF3B4 consensus motif. This was reflected by a much lower percentage of the T allele in the SF3B4 CLIP-seq data than in the RNA-seq data and increased usage of an upstream cryptic 3’ splice site for the TT genotype (Fig. 6c, d). Overall, our results show that ASPRIN can help pinpoint causal variants within a window of SNPs that are correlated with levels of alternative splicing and in high linkage disequilibrium with each other.

**Figure 6:**
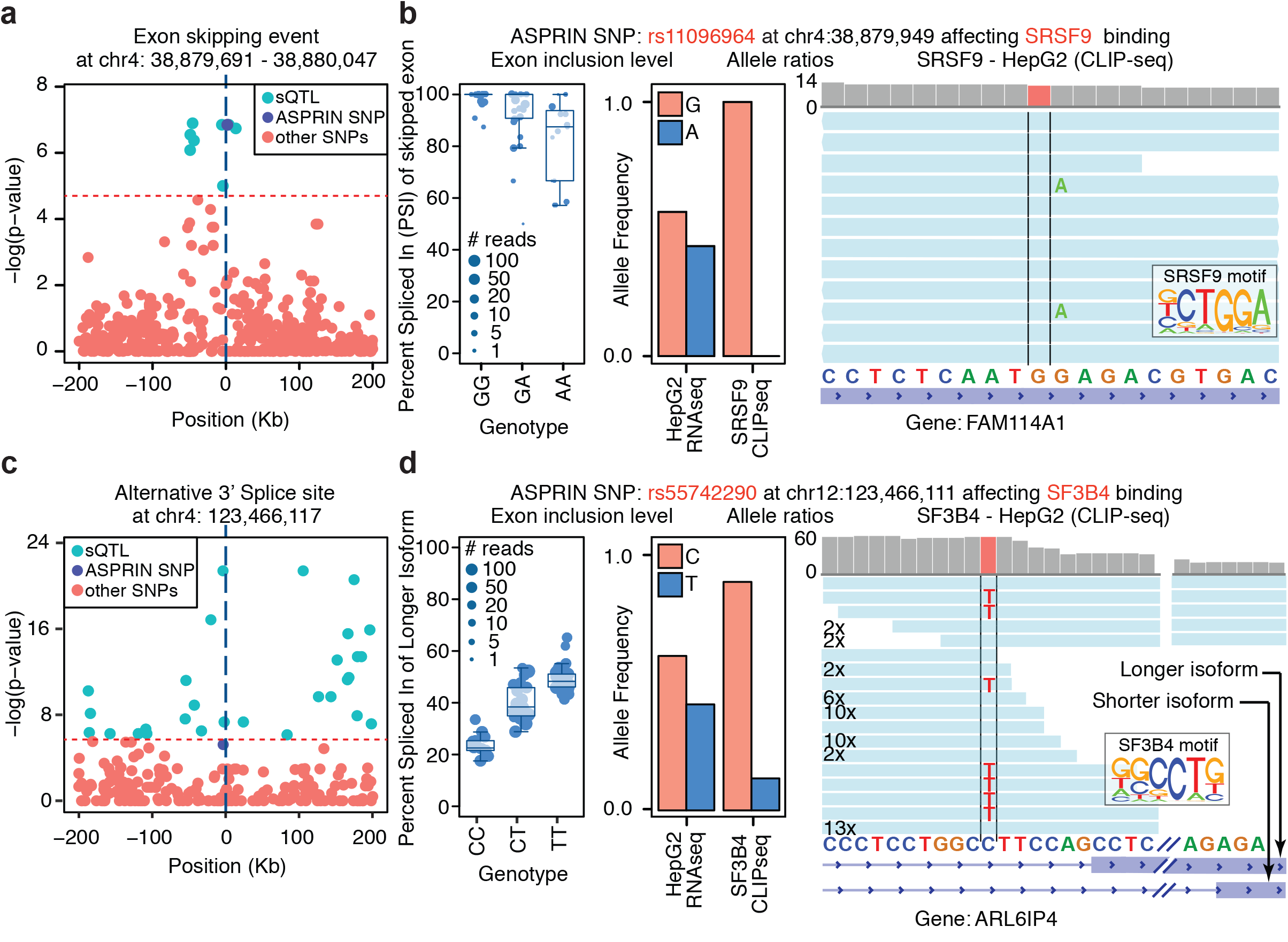
ASPRIN helps reveal causal variants affecting alternative splicing. (**a**) Distribution of GLiMMPS *p*-values around the exon skipping event in FAM114A1. For each SNP, the *p*-value indicates the significance of correlation between genotype and exon inclusion level within a 400-kb window centered on the splicing event. (**b**) Plots indicating the correlation of exon inclusion level with genotype for the ASPRIN SNP, differential binding of SRSF9 to the ASPRIN SNP that is in high LD with the GLiMMPS sQTL, and CLIP-seq allelic coverage on the ASPRIN SNP illustrating the effect of the SNP on the RBP consensus motif. (**c, d**) Similar plots are shown for a GLiMMPS sQTL involving alternative 3’ splice site usage in ARL6IP4, along with an ASPRIN SNP with differential binding of SF3B4 that is in high LD with the sQTL.

We further associated ASPRIN SNPs with genome-wide association study (GWAS) SNPs [28]. Specifically, we used the linkage disequilibrium (LD) map of a CEU population to calculate LD correlations between all ASPRIN SNPs and SNPs associated with diseases and traits in the NHGRI GWAS catalog [28]. Supplementary Tables 1 and 2 show all ASPRIN SNPs in high LD (r^2^ > 0.8) with a GWAS SNP in HepG2 and K562 cell lines respectively. These tables can be used by researchers to narrow down their search for candidate causal SNPs from GWAS signals of human traits or diseases.

## Discussion

We report ASPRIN, a computational tool for identifying genetic variants that may affect RBP-RNA interactions, by quantifying and contrasting the allelic ratios of heterozygous SNPs in CLIP-seq vs. RNA-seq data. Unlike previous work that relied on short RBP consensus motifs [15, 16], ASPRIN adopts a data-driven approach to directly observe the allelic preference of RBPs in CLIP-seq data, using matching RNA-seq from the same cell type as the control. Our comprehensive ASPRIN analysis of 166 RBPs in two ENCODE cell lines identified 55,646 candidate allele-specific protein-RNA interaction events. These events may provide valuable information for interpreting causal signals underlying human transcriptomic variation and phenotypic diversity. Of note, recent population transcriptomic studies (such as the GTEx project [29]) have revealed widespread genetic variation of gene expression and RNA processing in human populations, but identifying the causal SNPs underlying such regulatory variation remains difficult. The ASPRIN analysis provides an independent source of information that may assist the fine mapping of SNPs associated with gene expression levels or RNA processing patterns. In this work, we present two example cases in which the ASPRIN analysis reveals the likely causal variant responsible for splicing QTLs in the human liver. Future studies integrating other layers of RNA regulatory processes may reveal ASPRIN SNPs that causally impact other aspects of RNA processing and metabolism in human cells.

## Methods

### Calling variants from RNA-seq data

The total RNA-seq data for HepG2 whole-cell preparations from two different labs (ENCSR468ION and ENCSR181ZGR), a HepG2 cytosolic fraction (ENCSR862HPO), a HepG2 nuclear fraction (ENCSR061SFU), K562 whole-cell preparations from two different labs (ENCSR000AEN and ENCSR885DVH), a K562 cytosolic fraction (ENCSR860DWK), and a K562 nuclear fraction (ENCSR040YBR) were downloaded from the ENCODE website (https://www.encodeproject.org/).

The GATK Best Practices workflow for SNP and indel calling on RNA-seq data was used with minor modifications [18]. Briefly, the data sets were mapped using STAR v.2.5.2a [30], and total RNA-seq data from all fractions and all labs were merged to make one large RNA-seq data set for each cell line. The rest of the pipeline included adding read groups, sorting, marking duplicates, and creating the index using Picard tools v1.134 [31], followed by splitting and trimming, subsequent reassignment of mapping qualities, indel realignment, base recalibration, and finally, calling variants using GATK pipeline [18] (command lines in Supplementary Information). Mapping and variant calling statistics are given in Supplementary Table 3.

### Filtering SNPs

To remove false positive SNPs due to sequencing errors, alignment artifacts, and RNA editing events, only heterozygous variants in the RNA-seq data that matched known SNPs in dbSNP v150 [19] were kept. In this analysis dbSNP v150 was downloaded from: ftp://ftp.ncbi.nih.gov/snp/organisms/human_9606/VCF/All_20170710.vcf.gz. Potential RNA editing events were labeled and removed by intersecting our called heterozygous variants with the RADAR (Rigorously Annotated Database of A-to-I RNA editing) RNA editing database [20]. RADAR database version 2 was downloaded from: http://lilab.stanford.edu/GokulR/database/Human_AG_all_hg19_v2.txt.

### eCLIP data analysis

For analyzing ENCODE eCLIP data, the Standard Operating Procedure (SOP) published on the ENCODE website (https://www.encodeproject.org/) was followed. In brief: (1) adaptors were trimmed using cutadapt v1.10 [32]; (2) a second round of adaptor cutting was performed to control for double ligation events; (3) the resulting reads were mapped to the human-specific version of Repbase [33] using STAR 2.5.2a [30] to remove repetitive elements and other repetitive reads, as well as to control for spurious artifacts from rRNA. Repbase was downloaded from: http://www.girinst.org/downloads/; (4) Reads mapped to repetitive regions were filtered out of the resulting output from STAR; (5) PCR duplicates were further removed using random-mers, which are provided in the names of the reads; (6) Second (paired-end) reads were used to perform peak-calling using Piranha [23], with a bin size of 1nt. We considered significant peaks to be those that had a corrected *p*-value of less than 0.01 (command lines in Supplementary Information). Mapping and peak calling statistics for RBPs in HepG2 and K562 cell lines are given in Supplementary Tables 4 and 5 respectively.

### ASPRIN allelic ratio test

For each RBP, ASPRIN counts the number of reads that cover each allele in the CLIP-seq and RNA-seq data sets and forms a contingency table with: (1) the number of reads covering the reference allele in CLIP-seq; (2) the number of reads covering the alternative allele in CLIP-seq; (3) the number of reads covering the reference allele in RNA-seq; and (4) the number of reads covering the alternative allele in RNA-seq. The result of Fisher’s exact test for each SNP shows whether a particular SNP is significantly differentially bound by an RBP. For each RBP, ASPRIN *p*-values are corrected for multiple hypothesis testing using the Benjamini-Hochberg method and reported SNPs with *q*-value < 0.1 as significantly differentially bound, or “ASPRIN SNPs”.

### Assessing the robustness of ASPRIN

To measure the error associated with the used variant filtering method, RNA-seq data sets for the GM12878 cell line were downloaded from SRA (SRR307897 and SRR307898) and the complete genotype for this cell line from the 1000G website [21]. We performed variant calling as described above and intersected the set of called variants with 1000G SNPs, dbSNP, and RADAR.

To investigate the choice of RNA-seq protocol and how it may affect the power of ASPRIN, in addition to the total RNA-seq data, we also downloaded polyA+ mRNA-seq data for the same cell lines, fractions, and laboratories: HepG2 whole-cell preparations from two different labs (ENCSR985KAT and ENCSR561FEE), a HepG2 cytosolic fraction (ENCSR931WGT), a HepG2 nuclear fraction (ENCSR058OSL), K562 whole-cell preparations from two different labs (ENCSR000AEO and ENCSR545DKY), a K562 cytosolic fraction (ENCSR384ZXD), and a K562 nuclear fraction (ENCSR530NHO). To normalize for read number and length, we sampled *n* number of reads from all of these data sets, ten times, where *n* was the minimum number of reads among these data sets. The RNA-seq libraries that had 100-nucleotide reads (from Brenton Graveley’s lab) were also truncated to 50 nucleotides, to have the same read length as the RNA-seq libraries with 50-nucleotide reads (from Eric Lecuyer’s lab). We then called variants from all these data sets and compared the number of called variants and the regions in which these variants were located. We also ran the ASPRIN pipeline on all CLIP data sets with these ten subsampled RNA-seq data sets using only cytosolic polyA+ mRNA-seq and nuclear total RNA-seq, to compare the number of ASPRIN SNPs that can be called using these two distinct RNA-seq sets representing different RNA species and subcellular fractions.

To investigate the cross-linking bias and its potential effects on our analysis, for any ASPRIN SNP that was associated with at least one of the 75 RBPs in the HepG2 cell line, we counted for how many RBPs this SNP was: (1) called significant with preference for the reference allele; (2) called significant with preference for the alternative allele; (3) not called significant; and (4) not present in enough reads to pass the filters for the ASPRIN analysis.

### ASPRIN SNP enrichment or depletion in genomic regions

We measured the enrichment of ASPRIN SNPs in different genomic regions using Fisher’s exact test. For RBP *x* and region *r*, we counted (1) the number of ASPRIN SNPs for *x* in *r*, and (2) the rest of ASPRIN SNPs for *x*. In addition, for the background, we counted (3) the number of ASPRIN SNPs in *r* for the rest of the RBPs and (4) the number of ASPRIN SNPs in any region except *r* for the rest of the RBPs. Then, we used Fisher’s exact test to measure the significance of enrichment or depletion of ASPRIN SNPs in region *r* for RBP x compared to the average expectation.

### Measuring RBP sequence specificity

We determined the sequence specificity of RBPs as the information content of the motif obtained by *de novo* motif discovery in the high-quality binding sites as defined by the Piranha peak caller [23]. For each RBP, peaks output was obtained using Piranha. Then the genomic region (intron, 5' UTR, coding segment, 3' UTR, noncoding RNA, and intergenic sequence) containing each peak was assigned to them. All peaks in noncoding or intergenic regions were filtered out and the highest peak in each gene was selected as the representative peak of that RBP binding to the gene. Finally, top 1000 peaks based on the corrected *p*-value reported by Piranha were selected as the set of high-quality peaks. Zagros [8] was then used for *de novo* motif discovery, using sequence and secondary structure information. The parameters were window size 6 and top 10 motifs (-w 6 -n 10), and we selected the top motif reported by Zagros as the discovered motif for that RBP. Information content for each RBP consensus motif is obtained by taking the average information content over all positions within the consensus sequence and for each position defined by Shannon's entropy. RBPs with consensus sequences that had more information content were considered to have higher sequence specificity.

### Motif enrichment analysis

Motif enrichment analysis was done using the STORM software [26]. As described above the top 6-nucleotide motif discovered by Zagros in top 1000 peaks for each RBP was used as the consensus motif for that RBP. STORM can use the motif position weight matrix output from Zagros directly and calculate the enrichment of that motif in the set of input sequences.

For each SNP, a sequence of 11 nucleotides centered at the SNP (windows containing all 6-mer positions in the genome that include the SNP) was extracted. Then for each sequence we flipped the center nucleotide, the SNP, to the alternative allele. Therefore, for each RBP, two sets of sequences are formed, that are pairwise identical, except for the center position that contains two alleles of the SNP. One set contains the alleles with low affinity binding and the other contains the alleles with high affinity binding. Then STORM was run using the corresponding consensus motif for each RBP in two sets of sequences for the said RBP to assess the difference in motif score. Parameters for STORM can be set in a way to find the top occurrence of a motif per sequence (-n 1 -q) in single stranded mode (-S) for RNA (command lines in Supplementary Information). For each RBP we only considered SNPs that have positive scores in both high and low binding affinity sequences to filter out SNPs occurring outside the binding site. For each SNP the maximum motif score among all 6 possible windows on the high binding affinity sequence and its corresponding motif score in the low binding affinity sequence were selected to produce the boxplots of motif scores for each RBP in each position of the motif. We also defined a motif impact score for each RBP and its associated ASPRIN SNP set as the maximum difference in average motif score between the two alleles with high versus low binding affinity in the window of 6 nucleotides overlapping the ASPRIN SNP (Supplementary Fig. 7).

### sQTL analysis

To demonstrate the utility of ASPRIN in finding relevant SNPs that may cause changes in splicing, we analyzed ASPRIN SNPs in HepG2 cell line and sQTLs calculated from population-scale RNA-seq data in liver as part of the GTEx consortium [29]. RNA-seq and genotype data of liver tissues from 71 individuals (GTEx v6) were downloaded, mapped to the hg19 genome and Percent Spliced In (Psi) values were calculated for each splicing event in each individual. We selected events requiring the condition *Max(Psi) – Min(Psi) > 0.1* over all individuals. Then, for each splicing event, GLiMMPS [27] was run on SNPs within a 400-kb window centered on the splicing event. The false discovery rate (FDR) was estimated using a permutation procedure to obtain the null hypothesis. We performed this permutation ten times, recorded the minimum *p*-value for each site over all *cis* SNPs in each permutation, and used this set of *p*-values as the empirical null distribution. Using an FDR threshold of 10%, we calculated the *p*-value cutoff *t* such that *P*(*p*_0_ < *t*)/*P*(*p*_1_ < *t*) = 0.1, where *P*(*p*_0_ < *t*) is the fraction of expected *p*-values from the null distribution less than *t* and *P*(*p*_1_ < *t*) is the fraction of observed *p*-values less than *t* from the real data. For each splicing event, the sQTLs are defined as the SNPs that have *p*-values less than the cutoff. The linkage disequilibrium (LD) with all the ASPRIN SNPs was calculated and used for selecting only the exons that had sQTLs in high LD with ASPRIN SNPs (r^2^ > 0.8). The LD map was created using a CEU population [28]. Exons for cases in which the ASPRIN SNP is near the exon were further filtered with the criteria that the ASPRIN SNP is within a window of 500 nucleotides around the alternative splicing event. The windows were defined for each alternative splicing event as follows: (1) skipped exon: 500 nucleotides into the introns on each side of the skipped exon; (2) mutually exclusive exons: 500 nucleotides into the introns on each side of two mutually exclusive exons; (3,4) alternative 5' or 3' splice sites: 500 nucleotides into the introns on each side of the longer exon; and (5) intron retention: 500 nucleotides into the exons on each side of the retained intron. The numbers of exons in each type of alternative splicing event that pass the filters are given in Supplementary Table 6.

### GWAS signals

23,444 GWAS SNPs with *p*-values < 10^-5^ were downloaded from the NHGRI GWAS catalog [28] and PLINK v1.08p [34] was used to calculate the LD between ASPRIN SNPs and GWAS SNPs on the LD map that was created using a CEU population [28]. SNPs in high LD (r^2^ > 0.8) with GWAS SNPs were reported as GWAS-correlated ASPRIN SNPs.

### Software availability

The ASPRIN source code is available under GNU General Public License version 3.0 and can be downloaded from GitHub: https://github.com/Xinglab/ASPRIN.

## Acknowledgements

The authors thank the ENCODE Consortium and the ENCODE production laboratories for generating the eCLIP and RNA-seq data. This work was partly supported by the NIH T32 Tumor Cell Biology Training Grant (T32CA009056).

## Author Contributions

E.B.S. and Y.X. designed the methods, implemented the software, performed the analyses, and wrote the manuscript.

## Competing financial interests

The authors declare no competing financial interests.

